# Sustained Interleukin-1β overexpression exacerbates Tau pathology in a murine tauopathy model via cyclooxygenase-1

**DOI:** 10.1101/2021.02.10.430570

**Authors:** Simantini Ghosh, Solomon S. Shaftel, Stephanos Kyrkanides, John A. Olschowka, M. Kerry O’Banion

**Affiliations:** Department of Neurobiology and Anatomy, University of Rochester School of Medicine and Dentistry, Rochester, New York; Health Sciences Center, Stony Brook University, Stony Brook, New York

**Keywords:** Alzheimer’s disease, JNPL3, Microglia, Neuroinflammation, SC560

## Abstract

Pathologic accumulation of abnormally phosphorylated tau in neurofibrillary tangles is a hallmark feature of Alzheimer’s disease and other tauopathies. Interleukin-1β ◻◻◻ −1β◻ is a major proinflammatory cytokine in the central nervous system that has been implicated in the pathogenesis of tauopathies as well as Alzheimer’s disease. To explore the role of chronic IL-1β overexpression in tauopathies *in vivo* we used an inducible model of IL-1β overexpression developed in our laboratory. The IL-1β (IL-1) mice bear a transcriptional stop flanked by LoxP elements upstream of a human IL-1β gene. Upon delivery of Cre, the IL-1 transgene is locally activated by excision of the stop sequence. The IL-1 mice were bred to JNPL3 (Tau) mice, which overexpress human tau with the P301L mutation. Expression of IL-1β was induced in the dentate gyrus of 8 to 8.5 month old progeny by stereotaxic injection of FIV-Cre. One and three months later, Tau/IL-1 mice demonstrated 2-4 fold increases in phospho-tau and glial activation. To attenuate IL-1β mediated inflammation, we reduced PGE_2_ production via pharmacological inhibition of cyclooxygenase-1 (COX-1) with SC560 in Tau/IL-1 mice, and observed significant reductions in phospho-tau pathology and microglial activation. Further, we found upregulation in active forms of p38MAPK, which was significantly reduced in mice receiving SC560 treatment. Our results demonstrate that IL-1β has a direct exacerbating effect on tau pathology *in vivo*, and inhibiting COX-1 can reverse this. COX-1 inhibition can therefore serve as a valuable therapeutic strategy for tauopathies with an advanced inflammatory component.

## 1. Introduction

Tauopathies are a class of neurodegenerative disorders that result from anomalies in the microtubule associated protein tau (MAPT) gene, and demonstrate abnormal phosphorylation and aggregation of the axonal protein tau in the brain (Ballatore et al. 2007). Apart from Alzheimer’s disease (AD), tauopathies include progressive supranuclear palsy (PSP), corticobasal degeneration (CBD) and a subset of frontotemporal dementia (FTLD-tau) (Noble et al. 2013). These are swiftly progressing disorders with an average age of onset in the mid-fifties. Within this age group they are approximately as common as AD, but with quicker fatality (Roberson 2012). Persistent neuroinflammation is a common feature of these disorders, with evidence of active gliosis and upregulation of several neuroinflammatory chemokines and cytokines (Boxer et al. 2013a). Microglial activation correlates with tau pathology in CBD and PSP (Ishizawa and Dickson 2001). However, the exact role of neuroinflammation in tau pathogenesis is poorly understood.

Interleukin-1β ◻◻◻◻◻ β◻ is a pleiotropic, pro-inflammatory cytokine that is upregulated in response to CNS injury or chronic neurodegeneration, and has important roles in regulating the severity and length of the neuroinflammatory response (Tuppo and Arias 2005;Heneka and O’Banion 2007). IL-1 is prominently upregulated in Alzheimer’s disease (AD), the most common form of tauopathy, and has been investigated at length *in vitro* and *in vivo* in both acute and chronic injury models (Shaftel et al. 2008). The interest in IL-1 and its role in neurodegeneration were spurred by seminal observations that IL-1α positive glial cells were associated with senile plaques, dystrophic neurites and neurofibrillary tangles in AD brains, and that this association correlated with the severity of these pathological features (Sheng et al. 1995, 1997b, a). The interrelationship of IL-1 and amyloid has been the focus of intense research in the last decade, with contradictory results from *in vitro* and *in vivo* experiments (Goldgaber et al. 1989;Araujo and Cotman 1992;Gray and Patel 1993;Brugg et al. 1995;Barger and Harmon 1997;Griffin et al. 1998;Benzing et al. 1999;Shaftel et al. 2007a;Shaftel et al. 2007b;Matousek et al. 2012). However, the contribution of IL-1β to tau pathology under chronic neurodegenerative conditions is not well studied. One study reports an upregulation in IL-1β in neuronal and glial cells of a patient with Frontotemporal Dementia (FTD) (Bellucci et al. 2011). IL-1β treatment exacerbates tau phosphorylation through p38MAPK and downregulates synaptophysin in neuronal-microglial co-cultures (Li et al. 2003b). Rat brains impregnated with slow releasing IL-1 pellets demonstrate elevated p38MAPK mRNA, suggesting a potential upregulation in tau phosphorylation (Sheng et al. 2001).

We have previously described a role of chronic IL-1β overexpression in exacerbating tau pathology in the 3×TgAD mice (Ghosh et al. 2013). However, the 3×TgAD mice demonstrate both amyloid and tau pathology, hence a potential confound arises if the changes in tau pathology are secondary to amyloid modulation in these mice. Therefore, to understand the interrelationship of tau and IL-1β more thoroughly, we used an inducible model of sustained IL-1β overexpression developed in our laboratory (IL-1β^XAT^ mice; (Shaftel et al. 2007a;Shaftel et al. 2007b;Hein et al. 2010)) to overexpress transgenic human IL-1β in JNPL3 mice, a well characterized murine model of tauopathy that overexpresses the 4R0N isoform of the human tau transgene with the P301L mutation under a mouse prion promoter (Lewis et al. 2000).

Based on existing literature and our own data, we hypothesized that chronic upregulation of IL-1β would exacerbate phospho-tau pathology *in vivo*. In accordance with this hypothesis, we report elevated tau pathology following sustained overexpression of IL-1β in JNPL3/IL-1β^XAT^ mice (Tau/IL-1). We further investigated if attenuating neuroinflammation in these mice would reverse increased tau pathology. A prominent inflammatory mediator upregulated by IL-1β in the CNS is Prostaglandin E_2_ (PGE_2_). PGE_2_ is upregulated five fold in AD and can directly induce deficits in hippocampal learning in animals (Montine et al. 1999;Hein and O’Banion 2009). In the IL-1β^XAT^ mice, PGE_2_ production depended on Cyclooxygenase-1 (COX-1) activity (Matousek et al. 2010). Therefore, we treated Tau/IL-1 mice and appropriate controls with SC560, a selective pharmacological inhibitor of COX-1. Following a week of SC560 treatment, Tau/IL-1 mice demonstrated a marked reduction in phospho-tau pathology despite activation of the IL-1β transgene for 4 weeks. In addition we obtained evidence that the tau kinases, p38MAPK and GSK3β, were activated after induction of IL-1β in the Tau/IL-1 mice, and that this was blunted by SC560 treatment. In summary, our findings suggest that IL-1β is an important regulator of tau pathology *in vivo*, and interrupting the IL-1β mediated inflammatory pathways via COX-1 inhibition can serve as a valuable therapeutic strategy for tauopathies with an advanced inflammatory component.

## 2. Materials and Methods

### 2.1. Transgenic mice

All animal procedures were reviewed and approved by the University Committee on Animal Resources of the University of Rochester Medical Center for compliance with federal regulations prior to the initiation of the study. Two lines of transgenic mice were used in the present study. The construction and characterization of the IL-1β^XAT^ mice on a C57/BL6 background has been described previously (Shaftel et al. 2007a;Shaftel et al. 2007b). The JNPL3 mice (Lewis et al. 2000) express human tau with the P301L mutation under the mouse prion promoter, and develop neurofibrillary pathology starting in the spinal cord that spreads through the brain stem to the forebrain with age. Hemizygous male JNPL3 mice on a mixed B6D2F1/SW background (Line 001638-T) were purchased from Taconic Farms, Inc. and set up as breeders with hemizygous IL-1β^XAT^ females. One JNPL3 male breeder was set up with a Wild-Type (WT) female on a C57/BL6 background to provide mice for the time course experiment. These crosses were carried out in accord with an agreement between the University of Rochester and Taconic Farms, Inc. F1 progeny from these crosses were used for all experiments in this study with littermate controls.

For the time-course experiment we used 3-4 mice of each genotype (Tau and WT) for 7, 9 and 11-month time points each (9 WT and 13 Tau mice, 22 animals total). For the IL-1 overexpression experiment, we used the following genotypes for each of the 1 and 3 month time points: JNPL3/IL-1β^XAT^ (Tau/IL-1), JNPL3 (Tau), IL-1β^XAT^ (IL-1), and WT. 6-11 mice were used for each genotype at each time-point for immunohistochemical analysis (64 mice total). Gender balanced cohorts were used for these experiments. For the SC560 study, Tau (15; 5M, 10F) and Tau/IL-1 (15 (11M, 4F) mice were treated with SC560 or vehicle (30 mice total). 3 Tau male and 3 Tau/IL-1 female animals assigned to the SC560 group did not survive the three week post surgery period before SC560 treatment began.

### 2.2. FIV-Cre

The construction and packaging of the Feline Immunodeficiency Virus has been described previously (Lai et al., 2006). Briefly, the FIV-Cre virus encodes a modified version of Cre recombinase with a nuclear localization signal and a V5 epitope tag under the control of a Cytomegalovirus promoter. FIV-Cre was packaged to a final titer of approximately 1 × 10^7^ viral particles/ml.

### 2.3. Stereotaxic activation of the IL-1β transgene

Mice were anesthetized with 1.75% isoflurane in 30% oxygen and 70% nitrogen and secured in a Kopf stereotaxic apparatus using ear bars. Hair was removed with a commercially available hair removal lotion and the exposed skin was cleaned thoroughly with an alcohol swab to remove stray hairs. The scalp was thoroughly rubbed with betadine. An incision was made on the top of the skull, a 0.5 mm burr hole drilled in the skull at 1.8mm caudal and 1.8 mm horizontal from bregma, and a 33 GA needle connected to a 10 μl syringe (Hamilton Company) pre-loaded with virus was lowered 1.8 mm from the brain surface over 2 minutes. A Micro-1 microsyringe pump controller (World Precision Instruments) was used to inject 1.5 μl of virus at a constant rate over 10 minutes. Following 5 minutes to allow viral diffusion (about 1.5 × 10^4^ infectious viral particles), the needle was raised slowly over 2 minutes, the burr hole was sealed with Ethicon bone wax and soft tissues sealed using 3M Vetbond surgical glue (3M Animal Care Products). For the IL-1β overexpression study, all mice received a unilateral injection of FIV-Cre in the dentate gyrus to control for injuries resulting from the surgical procedure and injection of the FIV-Cre construct. For the SC560 study all animals received bilateral injections of FIV-Cre in their dentate gyrus.

### 2.4. SC560 treatment

SC560 has been reported to have an IC_50_ of 9 nM for COX-1 versus 6.3 ◻ M for COX-2 using recombinant enzymes in a cell free system (Smith et al. 1998). There is one study suggesting that SC560 is not COX-1 selective in primary neurons (Brenneis et al. 2006), however, a more recent report using equine whole blood again demonstrates SC560 being COX-1 selective (Cuniberti et al. 2012). The field lacks substantial evidence of a mechanistic comparison of these differential effects. Our choice of SC560 was propelled by the relatively well-documented history of SC560 usage as a selective COX-1 inhibitor (Pepicelli et al. 2005;Choi et al. 2008;Garcia-Bueno et al. 2009;Aid and Bosetti 2011;Choi et al. 2013;Griffin et al. 2013) and our own experience with SC560 (Moore et al. 2005;Matousek et al. 2010). Further, SC560 has been shown to exert almost no effect on COX-2 activity in rats compared to COX-1 (Smith et al. 1998). Other studies conducted on COX-1 selectivity of SC560 have been reviewed (Perrone et al. 2010). Tau/IL-1 and Tau mice received bilateral stereotaxic injections in the dentate gyrus with FIV-Cre as described above. Three weeks following injection, mice received daily intraperitoneal (i.p.) injections of 10 mg/kg SC560 (Cayman Chemicals) solubilized in vehicle (40% DMSO in 0.1M PBS pH 7.4; 9 Tau/IL-1, 8 Tau) or vehicle alone (6 Tau/IL-1, 7 Tau) for 7 days. This dosing regimen was selected based on our previous observations in IL-1β^XAT^ mice (Matousek et al. 2010) and previous literature (Blais et al. 2005;Garcia-Bueno et al. 2009). All mice were sacrificed 4 hours after the final drug injection. Hemibrains were collected as described below for immunohistochemical and molecular analysis.

### 2.5. qPCR

Mice were anesthetized with ketamine (i.p. 100mg/kg) and xylazine (i.p. 10mg/kg) and perfused with 0.15 M phosphate buffer (PB) containing 2IU heparin/ml, 0.5% w/v sodium nitrite, and 1 ◻ M indomethacin. Brains were removed and hippocampi microdissected, rapidly frozen in ice- cold isopentane, and stored at −80°C. Hippocampal tissue was homogenized using an Omni Tissue Homogenizer (Omni) and RNA was isolated using the Trizol (Invitrogen) extraction protocol. PCR reactions were carried out in a final volume of 20 ◻ l reactions containing iQ Supermix (Bio-Rad), 5 nM FITC dye, and custom designed primers and probes (Invitrogen). Samples were denatured at 95°C for 3 minutes, followed by 30 cycles of denaturing at 95°C for 60 sec, annealing at 60°C for 60 sec and extension at 72°C for 60 sec. The GAPDH housekeeping gene was used to normalize determinations of mRNA abundance. Sequences used were as follows: (from 5’ to 3’; F=forward, R=reverse primer, and P=probe): hIL-1 transgene, F: ctcttcgcggtctttccagt R: tggaccagacatcaccaagc P: acgatctgcgcacctgtacgat; GAPDH, F: cccaatgtgtccgtcgtg, R: cctgcttcaccaccttcttg, P: tgtcatcatacttggcaggtttctccagg.

### 2.6. Immunohistochemistry

Mice were deeply anesthetized with a mixture of ketamine (i.p 100 mg/kg) and xylazine (i.p 10 mg/kg), perfused intracardially with 0.15M Phosphate Buffered Saline (PBS) containing 0.5% sodium nitrite (weight/volume) and 2 IU heparin/ml, followed by 4% ice cold paraformaldehyde (PFA), pH 7.2 in 0.15 M PBS. For the IL-1β overexpression study, brains were collected, fixed for 2 additional hours in 4% PFA (pH 7.2) at 4°C, equilibrated in 30% sucrose in PBS overnight, frozen in cold isopentane and stored at −80°C. For the SC560 study, hemibrains were collected from mice intracardially perfused with 0.15 PBS containing 1 μM indomethacin, 0.5% sodium nitrite (weight/volume) and 2 IU heparin/ml, and post fixed overnight in 4% ice cold PFA, pH 7.2 in 0.15 M PBS. They were further equilibrated in 30% sucrose in PBS overnight, frozen in cold isopentane and stored at −80°C. All brains were cryosectioned into 30 μm sections on a −25°C freezing stage microtome and free floating sections were stored in a cryoprotectant solution until assayed. For immunohistochemical protocols sections were washed to remove cryoprotectant, incubated in 3% H_2_O_2_, washed, blocked with 10% normal goat serum and incubated in primary antibody for 48 h. They were then washed and incubated with biotinylated secondary antibody for 2 h at room temperature and developed with Elite ABC kit and 3,3-diaminobenzidine (Vector laboratories). Sections were mounted, cleared and cover-slipped in DPX (VWR). For biotinylated primary antibodies the secondary antibody incubation was omitted. Primary and secondary antibody dilutions: Iba-1 (Wako) 1:5000; AT180 and HT7 (biotinylated) (Thermo Scientific) 1:10; pT205 (Invitrogen) 1:5000; MHCII (Pharmingen) 1:6000; biotinylated goat anti-rabbit IgG (Vector) 1:2000; biotinylated goat anti-rat IgG (Vector) 1:2000.

### 2.7. Image acquisition and analysis

Photomicrographic images were captured by a Zeiss Axioplan IIi microscope equipped with a SPOT camera (Advanced Diagnostics) linked to a SPOT advanced software running on a 64 bit Microsoft Windows XP computer. For quantification purposes flat-field corrected images were captured with a 10X objective, and sections closest to the site of injection (<120 μm) in either direction were analyzed in NIH ImageJ v1.47 (http://rsbweb.nih.gov/ij/). The boundary of dentate gyrus was defined with the polygon tool, images were thresholded with suitable filters to detect optimal staining and eliminate background, converted to binaries and area fractions determined for each marker analyzed. For the IL-1β overexpression study, area fraction from each hemisphere was used to calculate an ipsilateral contralateral ratio (I/C ratio) for every section. I/C ratios were then averaged across sections analyzed (2-3) to obtain an I/C ratio per animal for each dentate gyrus and CA1. I/C ratios were averaged across animals to generate and compare mean I/C ratios for different genotypes. For the SC560 study, area fraction from every section was averaged across 2-3 sections to generate a mean area fraction from an animal. Values from animals were averaged to generate mean area fractions per genotype for the dentate gyrus and CA1. Representative images were captured with a 20X plan-apochromatic lens. Final images and layout were created using Adobe Photoshop CS6 and Adobe Illustrator CS6.

### 2.8. ELISA and EIA

Mice were deeply anesthetized with a mixture of ketamine (i.p 100 mg/kg) and xylazine (i.p 10 mg/kg), perfused intracardially with 0.15 M PBS containing 1 μM indomethacin, 0.5% sodium nitrite (weight/volume) and 2 IU heparin/ml. Hippocampi were quickly dissected, frozen in isopentane and stored at −80°C until further processing. Hippocampi were homogenized in Tper (Thermo Scientific; 50 mg/ml) with protease and phosphatase inhibitor cocktails (Calbiochem), vortexed and sonicated. Mouse IL-1β (mIL-1β) was determined according to kit instructions (R&D Systems) using hippocampal lysates in Tper. To extract PGE_2_, 700 μl of methanol was added to 300μl of hippocampal lysate in Tper, and spun at 13,000 g for 20 minutes at room temperature. Supernatants were evaporated and stored at −80°C. Evaporates were reconstituted with 300 μl kit buffer prior to analysis. PGE_2_ EIA was performed and data analyzed according to instructions provided by the kit (Cayman Chemicals).

### 2.9. Western Blot

Brains were collected, hippocampi dissected and frozen as described in the last section. Hippocampi were homogenized in Tper (Thermo Scientific; 50 mg/ml) with protease and phosphatase inhibitor cocktails (Calbiochem), vortexed and sonicated. Hippocampal lysates were diluted in 2x sample buffer (125 mM Tris-HCl; 4% SDS; 20% glycerol), and protein concentration was determined by a BCA assay (Thermo Scientific). Protein (15-20 μg/per lane for most blots) was electrophoresed on Tris-HCL polyacrylamide gels and transferred to nitrocellulose membranes (Bio-Rad) for 90 minutes at 4°C. After 1 hour in Western blocking reagent (Roche Diagnostics), membranes were incubated overnight with primary antibodies. After rinsing, blots were incubated with peroxidase-linked secondary antibodies (provided in Suprasignal West Dura Kit; Thermo Scientific), treated with the ECL substrate included in the kit and bands were visualized using Kodak Biomax XAR film. List of Primary Antibodies used: Tubulin (Calbiochem) 1:5,000; anti Tau (Dako) 1:10,000; AT180 (Thermo Scientific) 1:100; pT205 (Invitrogen) 1:1000; PHF1 (gift of Dr. Peter Davies) 1:100; and phospho Ser9 GSK3β, GSK3β, phospho p38 MAPK, and p38 MAPK (Cell Signaling) 1:1000.

### 2.10. Statistics

All Statistical comparisons were performed using Prism 5.0 (Graphpad Software). A p-value ≤ 0.05 was considered significant. Student’s *t-*test was employed when two group means were compared. Results where more than two group means were compared were analyzed with one-way or two-way Analysis of Variance (ANOVA) depending on the number of parameters. A Bonferroni or Tukey’s posttest was used to establish significance between individual groups in such instances.

## 3. Results

### 3.1. Chronic neuroinflammatory phenotype in the ipsilateral dentate gyrus of the Tau/IL-1 mice following one and three months of transgene activation

In this study we sought to characterize the effects of sustained IL-1β on tau pathology in transgenic mice harboring a tau mutation. Because we bred the Tau mice to IL-1 mice on a C57BL/6 background and used F1 progeny that were hemizygous for both Tau and IL-1, we altered the genetic background of this line for our study. To investigate whether this altered the pathological timeline in these animals and to establish an appropriate time point for our experiments, we characterized tau pathology and baseline microglial activation in 7, 9 and 11 month old Tau mice (Fig. *S1*, *S5*). We could detect transgenic tau expression and pT205 epitope phosphorylation at 7 through 11 months of age in the hippocampus of F1 Tau mice, especially in the dentate gyrus (Fig. *S1*). Littermate age matched WT controls did not immunostain for either of these markers. Based on this data we decided to induce IL-1β transgene expression in F1 Tau/IL-1 mice at 8 months of age and analyze the brains of these mice after one and three months of transgene expression at 9 and 11 months of age, respectively (Fig. *1*). To excisionally activate the *h*IL-1β transgene, 8-month-old Tau/IL-1 mice received a unilateral stereotaxic injection of FIV-Cre in the dentate gyrus of the hippocampus. To control for viral transduction and surgical procedures, three littermate control genotypes, Tau, IL-1 and WT received similar stereotaxic injections with FIV-Cre. This experimental design allowed us to use the contralateral dentate gyrus and hippocampus as an internal control for every animal. Two separate cohorts of mice were injected for assessment at one and three months following transgene activation. At both time points assessed, Tau/IL-1 mice demonstrated a marked upregulation in MHCII immunostaining in the ipsilateral dentate gyrus and hippocampus, suggesting a robust local neuroinflammatory response (Fig. *S2*). Similar MHCII staining was observed in IL-1 mice, but not in Tau or WT mice, which lack the hIL-1β transgene, one (Fig. *S3*) and three months (data not shown) following viral transduction.

**Fig. 1.**
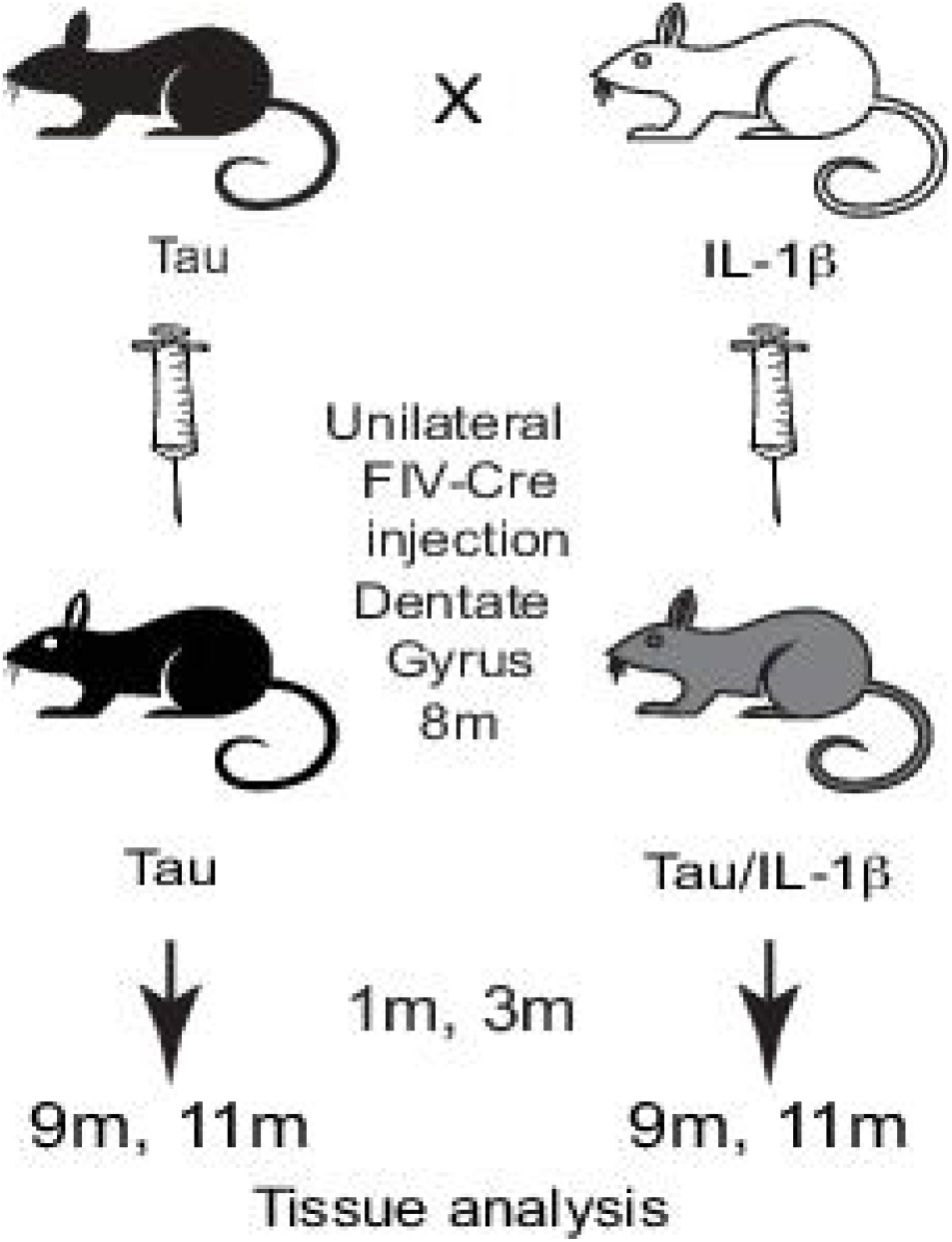
Research design. JNPL3 (Tau) mice were bred to IL-1β^XAT^ (IL-1) mice to obtain JNPL3/IL-1β^XAT^ (Tau/IL-1) mice and control JNPL3 (Tau) mice. All mice received a unilateral stereotaxic injection of FIV-Cre in the dentate gyrus of the hippocampus at 8 months of age to activate the IL-1β transgene and control for viral transduction. Cohorts of mice were sacrificed after 1 and 3 months for analysis. Two other control genotypes obtained from the breeders, wild type (WT) and IL-1β^XAT^ (IL-1), were used for this experiment but are not shown in this scheme.

### 3.2. Tau/IL-1 mice demonstrate exacerbated phospho-tau pathology and microglial activation in the ipsilateral dentate gyrus following one and three months of transgene activation

To study the effect of sustained IL-1β overexpression on tau pathology, we performed quantitative immunohistochemistry on sections <120 μm away from the FIV-Cre injection site one and three months following FIV-Cre transduction in all genotypes. One-month post activation, we found a robust upregulation in phospho-tau pathology, as shown by pT205 immunostaining in the ipsilateral dentate gyrus and overall hippocampus of the Tau/IL-1 mice compared to control Tau mice (Fig. 2*A*, *S4*). The exacerbated phospho-tau pathology persisted three months following FIV-Cre injection in the ipsilateral hippocampus of the Tau/IL-1 mice (Fig. *2B*). At any of the time points assessed, genotypes lacking the human tau transgene (IL-1 and WT) did not show any detectible staining for pT205. Quantification of the pT205 immunohistochemical data revealed 2-6 fold increases in pT205 immunostaining in the ipsilateral dentate gyrus and CA1 region of the hippocampus in the Tau/IL-1 mice (Fig. 3 *A,D*). To control for intra-animal variability, we present all data as Ipsilateral/Contralateral (I/C) ratios for analyzed parameters. Specifically, the I/C ratios for pT205 immunopositive area fraction were significantly upregulated approximately 3.7 and 6.4 fold in the dentate gyrus of the Tau/IL-1 mice compared to Tau mice (Mean I/C ratios for Tau mice: 0.99 and 1.04 respectively) at one and three months post transgene activation (Fig. 4*A*). In CA1 this increase was approximately 2.0 and 3.05 fold in the Tau/IL-1 mice compared to Tau mice (Mean I/C ratios for Tau mice: 1.0 and 1.1 respectively) at one and three months post transgene activation (Fig 3*D*). The data analyzed with a two-way ANOVA and Bonferroni post-hoc tests with genotype and duration of transgene expression as parameters, revealed an extremely significant effect of genotype on pT205 area fractions in both the dentate and the CA1 (F_(1,23)_=17.82, p=0.0003 for the dentate gyrus and F_(1,23)_=28.42,p<0.0001 for the CA1; n=5-11mice/group). We also observed upregulation in AT180 immunostaining and 3.67 fold increase in the I/C ratio of AT180 immunopositive area fraction of Tau/IL-1 mice compared to Tau mice one-month post transgene activation (p= 0.02;Fig *S3A,B*). To ascertain whether increased phospho-tau is not secondary to changes in human tau transgene expression, we immunostained sections <120 μm distant from the FIV-Cre injection site with HT7, an antibody that recognizes transgenic tau. In our analysis and quantification, mean I/C ratios of HT7 positive area fractions did not change significantly between any genotype at either time point in the dentate gyrus (Fig. 3*B;* F _(1,22)_= 0.88, p=0.35, n=5-11 mice/group) or in CA1 (Fig. 3*E;* F _(1,22)_= 2.85, p=0.10, n=5-11 mice/group). We also analyzed microglial activation in all animals at both time points with Iba1 immunohistochemistry. Iba1 immunoreactivity was markedly increased in the Tau/IL-1 mice at one and three months following transgene overexpression (Fig, 2*C,D*). Consistent with our previous observations, we found 2-3 fold elevated microglial activation in the Tau/IL-1 mice compared to Tau mice (I/C ratios 2.4 and 3.7 in the dentate of Tau/IL-1 mice vs. 0.96 and 0.98 in control Tau mice, and 3.0 in CA1 of Tau/IL-1 mice vs. 1.04 and 1.02 in control Tau mice at one and three months post transgene activation, respectively) (Fig. 3*C,F*). As with pT205, Iba1 immunohistochemical data analyzed by a two way ANOVA revealed an extremely significant effect of genotype at both time points analyzed for both the dentate gyrus (F _(1,22)_= 119.9, p<0.0001, n=5-11 mice/group) and the CA1 (F _(1,22)_= 47.33, p<0.0001, n=5-11 mice/group). In summary, we report increased immunostaining at the pT205 and AT180 phospho-tau epitopes in the Tau/IL-1 mice at the site of IL-1β overexpression, compared to the same brain regions in control Tau mice, with similar changes in microglial activation, but not transgenic tau. This reaffirms our findings from the 3xTgAD mouse model (Ghosh et al. 2013) and provides more evidence for a detrimental role played by sustained IL-1β overexpression in tau pathogenesis.

**Fig. 2.**
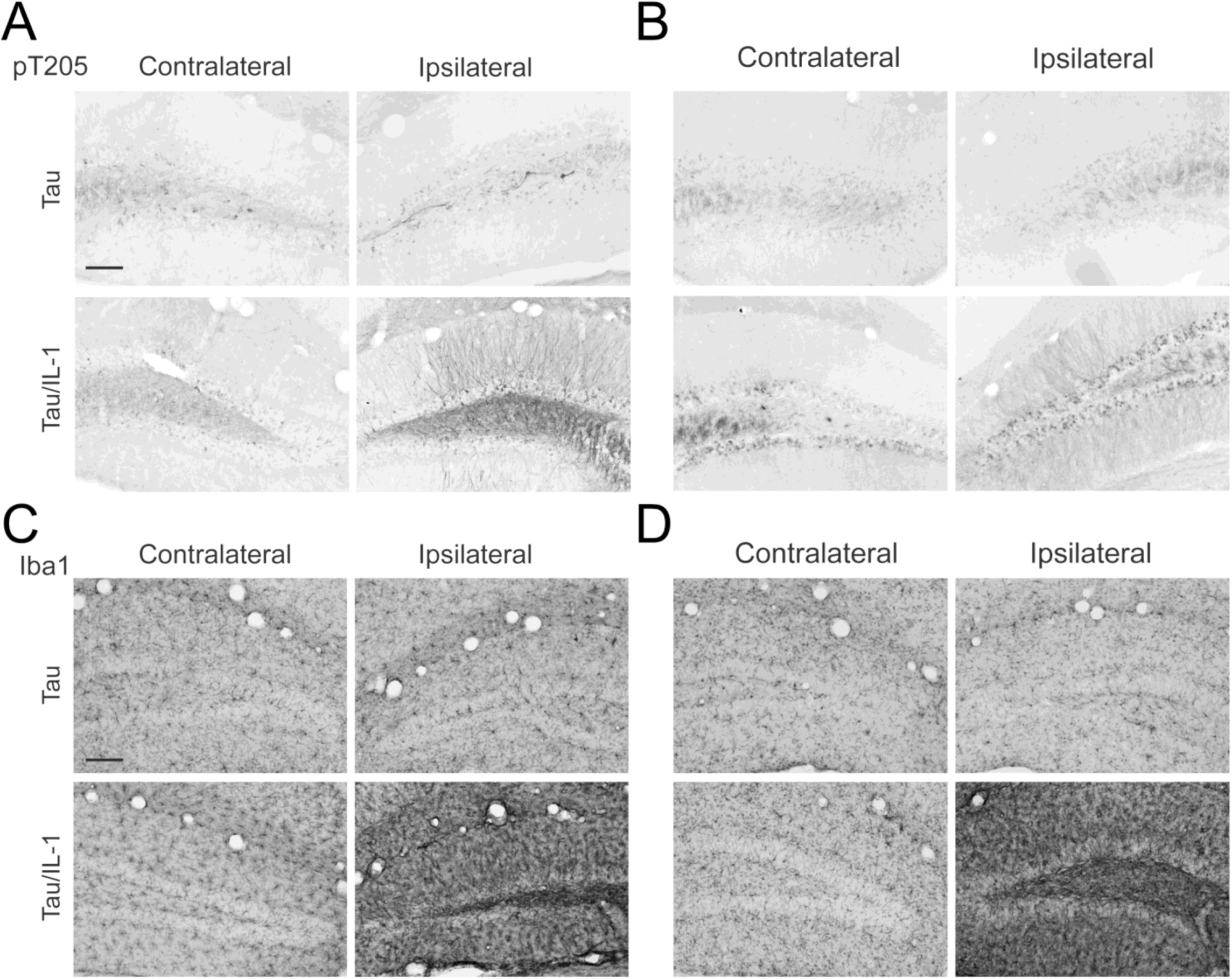
Tau/IL-1 mice demonstrate elevated tau phosphorylation and microglial activation but unaltered transgenic tau expression in the dentate gyrus after one and three months of transgene expression. Hippocampal sections of 9 and 11 month old Tau and Tau/IL-1 mice (one and three months of IL-1β overexpression, respectively) were probed with antibodies against phospho-tau (pT205) and a microglial marker (Iba1). Representative images of section stained with the phospho tau antibody pT205 (***A, B***) and the microglial marker Iba1 (***C, D***) showed increased immunostaining in the ipsilateral dentate gyrus of Tau/IL-1 mice at 9 (***A, C***) and 11 months (***B, D***) of age; Scale bars 100 μm.

**Fig. 3.**
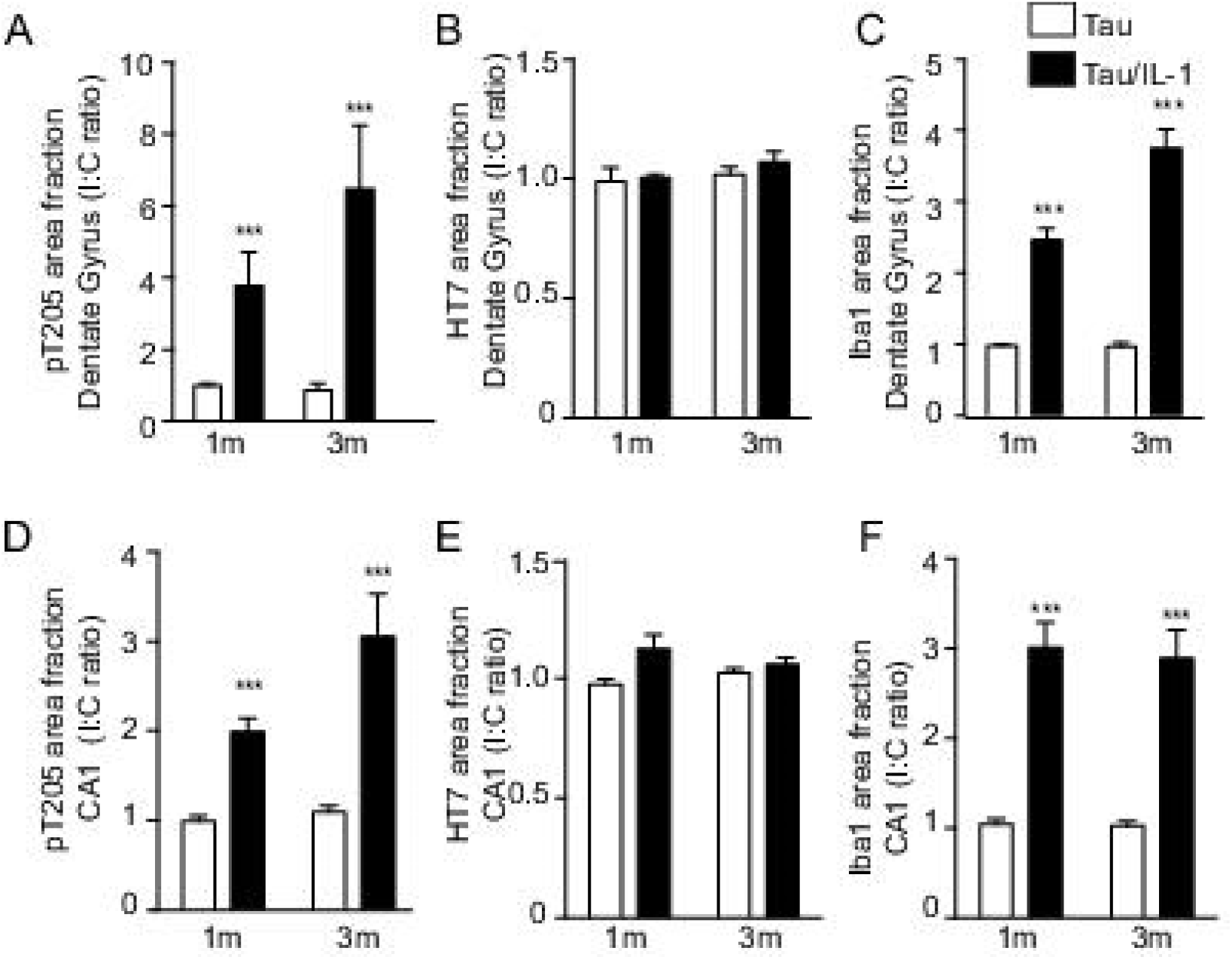
Tau/IL-1 mice demonstrate elevated tau phosphorylation and microglial activation but unaltered transgenic tau expression in the dentate gyrus after one and three months of transgene expression. Quantitative analyses of area fractions in the dentate gyrus and CA1 of 9 and 11 month old Tau (open bars) and Tau/IL-1 (black bars) mice are shown, following 1 and 3 months of transgene expression respectively. Immunopositive area fractions were quantified in the dentate gyrus (***A, B, C***) or in the CA1 (***D, E, F***) of the hippocampus. All numerical data is represented as mean Ipsilateral/Contralateral (I/C) ratio ± SEM per group. Tau/IL-1 mice demonstrated a 2-6 fold increase in pT205 immunostaining (***A, D***) and a 2-4 fold increase in Iba1 immunostaining (***C, F***) but not in HT7 immunostaining (***B, E***) in the ipsilateral hemisphere; n= 5-11 mice/group; data analyzed with two-way ANOVA with genotype and duration of transgene expression as parameters. Genotype had an extremely significant effect on pT205 and Iba1 immunostaining. *** p <0.001 (For detailed description of the quantitative data, please refer to the text).

**Fig. 4.**
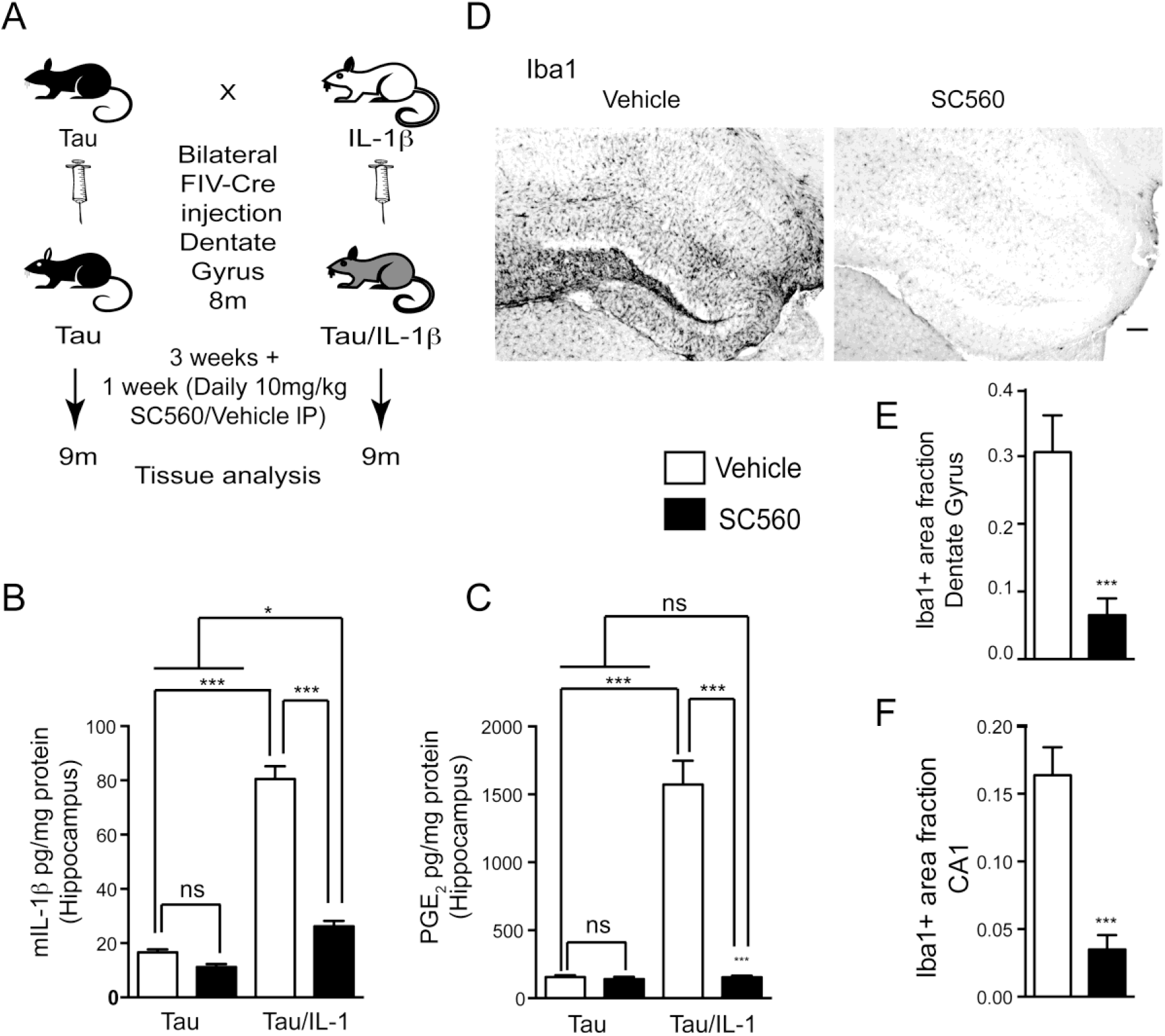
Attenuating inflammation with SC560 in Tau/IL-1 mice. ***(A)*** Research design: JNPL3 (Tau) mice were bred to IL-1β^XAT^ (IL-1) mice to obtain JNPL3/IL-1β^XAT^ (Tau/IL-1) mice and control JNPL3 (Tau) mice. Eight month old JNPL3 (Tau) and JNPL3/IL-1β^XAT^ (Tau/IL-1) mice received bilateral stereotaxic injections of FIV-Cre in the dentate gyrus of the hippocampus. Following three weeks of transgene expression, all mice received daily intraperitoneal injections of 10 mg/kg SC560 or vehicle for 7 days. All mice were killed for tissue analysis 4 hours after the final injection. The age of the mice at this end point was 9 months. ***(B-F)*** SC560 treatment reduced neuroinflammation in the hippocampus of 9 month old Tau/IL-1 mice overexpressing IL-1β for 4 weeks. Hippocampal ELISA measurements for mIL-1β ***(B)*** and EIA measurements for PGE_2_ ***(C)*** are shown for vehicle treated (open bars) and SC560 treated (black bars) Tau or Tau/IL-1 mice. ***(B)*** 4 weeks of transgene expression led to a significant upregulation in mIL-1β (Vehicle treated Tau vs. Tau/IL-1). One week of treatment with SC560 significantly reduced mIL-1β (Tau/IL-1 vehicle vs. SC560 treated). ***(C)*** Similar changes were seen in PGE_2_ levels following SC560 treatment. For detailed description of quantitative outcomes please refer to the text. All data are expressed as mean pg/mg protein ± SEM per group. Data analyzed by a one-way ANOVA with Tukey’s multiple comparison post-test for individual comparisons. n= 5-9/group. *** p<0.001; * p<0.05; ns: not significant. ***(D)*** Representative micrographs of sections from the dentate gyrus and CA1 of the hippocampus of 9-month-old Tau/IL-1 mice immunostained for microglial activation (Iba1) are shown. One week of treatment with SC560 reduced Iba1 immunostaining in the hippocampus of Tau/IL-1 mice despite overexpression of the IL-1β transgene for 4 weeks; Scale bars 100 μm. Quantification of Iba1 immunopositive area fraction in the dentate gyrus ***(E)*** and CA1 ***(F)*** of the hippocampus of Tau/IL-1 mice showed significant reduction in SC560 treated groups (black bars) compared to vehicle treated groups (open bars). All data are represented as mean Iba1 immunopositive area fraction ± SEM per group. Data analyzed by a Student’s t-test. n=6-9/group. *** p<0.001.

### 3.3. Pharmacological inhibition of COX-1 by SC560 attenuates the neuroinflammatory response in Tau/IL-1 mice despite activation of the IL-1β transgene

Next, we sought to investigate whether mitigating the neuroinflammatory response in Tau/IL-1 mice would reverse the exacerbation of phospho-tau pathology. To this end, we chose to target the COX-PGE_2_ pathway. PGE_2_ is a powerful inflammatory mediator that is upregulated by IL-1β in the IL-1 mice and can cause a functional deficit in contextual fear memory (Hein and O’Banion 2009). PGE_2_ production in the IL-1 mice is primarily mediated by COX-1 and not COX-2 (Matousek et al. 2010). We therefore set out to inhibit COX-1 dependent PGE_2_ production with the pharmacological inhibitor SC560 in Tau/IL-1 mice and evaluate phospho-tau pathology. For this experiment, we used Tau/IL-1 and littermate control Tau mice. The IL-1β transgene was activated bilaterally in the dentate gyrus of 8-month-old Tau/IL-1 mice. To control for viral transduction, control Tau mice received similar bilateral stereotaxic injections of FIV-Cre in the dentate gyrus. Three weeks following FIV-Cre injection, Tau/IL-1 and Tau mice were randomly assigned to vehicle or SC560 groups, and received a daily intraperitoneal injection of 10 mg/kg SC560 or vehicle for a week. All mice were sacrificed four hours after the final injection and their brains collected for tissue analysis (Fig 4*A*).

A distinct cohort of IL-1β overexpressor mice were utilized to analyze the expression of the hIL-1β transgene four weeks post injection with FIV-Cre, during the last week of which period mice were randomized to vehicle or SC560 groups. No change in the levels of transcript was detected between the vehicle and the SC560 groups (Fig. *S6*). Since IL-1β can sustain its own production to maintain a neuroinflammatory response, murine IL-1β (mIL-1β) ELISA measurements provide a reliable method to assess the extent of neuroinflammation in the IL-1 mice. Activation of the IL-1β transgene caused a marked upregulation in hippocampal mIL-1β (16.59 ± 1.13 pg/mg protein for vehicle treated Tau mice vs. 80.57± 4.67 pg/mg protein for vehicle treated Tau/IL-1 mice; p<0.001), which was significantly reduced in Tau/IL-1 mice that received SC560 treatment (26.24± 2.01 pg/mg protein; p <0.001; n =5-10/group; Data analyzed by a one-way ANOVA followed by a Tukey’s post-hoc test for individual comparisons; Fig 4*B*). Next, we investigated whether SC560 treatment lowered PGE_2_ production in the Tau/IL-1 mice. As measured by EIA, hippocampal PGE_2_ levels were dramatically reduced to control levels in Tau/IL-1 mice treated with SC560 (153.9± 12.15 pg/mg protein for SC560 treated Tau/IL-1 mice vs. 1573± 174.4 pg/mg protein for vehicle treated Tau/IL-1 mice; p <0.001). Vehicle and SC560 treated Tau mice showed no significant difference in hippocampal PGE_2_ content compared to SC560 treated Tau/IL-1 mice (155.7± 12.72 pg/mg protein and 141.6 ± 16.15 pg/mg protein vs. 153.9± 12.15 pg/mg protein respectively; n =5-10/group; Data analyzed by a one-way ANOVA followed by a Tukey’s post-hoc test for individual comparisons; Fig 4*C*). These findings of lowered PGE_2_ and mIL-1β levels indicate that SC560 treatment attenuates the neuroinflammatory response in Tau/IL-1 mice. To further confirm the attenuated neuroinflammatory response in SC560 treated mice, we performed Iba1 immunohistochemistry to characterize microglial activation. SC560 treated Tau/IL-1 mice showed a marked reduction in Iba1 immunoreactivity in the dentate gyrus as well as CA1 region of the hippocampus compared to vehicle treated Tau/IL-1 mice (Fig 4*D*). Quantification of Iba1 immunopositive area fractions demonstrated a ~5 fold reduction in the dentate gyrus and CA1 of SC560 treated Tau/IL-1 mice compared to vehicle treated Tau/IL-1 mice (mean Iba1 immunopositive area fractions in the dentate gyrus: 0.06 ± 0.02 for SC560 treated mice vs. 0.3 ± 0.05 for vehicle treated mice; Data analyzed by a Student’s t-test, p<0.001,t=4.36,df=14, n=7-9/group, Fig. 4*E*; Mean Iba1 immunopositive area fractions in the CA1: 0.03 ± 0.01 for SC560 treated mice vs. 0.16 ± 0.02 for vehicle treated mice; Data analyzed by a Student’s t-test, p<0.01,t=6.06,df=13, n=6-9/group; Fig. 4*F*).

### 3.4. Pharmacological inhibition of COX-1 by SC560 attenuates phospho-tau pathology in Tau/IL-1 mice despite activation of the IL-1β transgene

To test if SC560 treatment could reverse the exacerbation in phospho-tau pathology observed earlier, we investigated phospho-tau pathology by immunostaining for pT205 in the SC560 and vehicle treated Tau/IL-1 mice. We observed a significant decline in pT205 immunostaining in the dentate gyrus and CA1 region of the hippocampus in SC560-treated compared to the vehicle-treated Tau/IL-1 mice (Fig. 5*A*; Mean pT205 immunopositive area fractions in the dentate gyrus: 0.09 ± 0.01 for SC560 treated mice vs. 0.23 ± 0.02 for vehicle treated mice; Data analyzed by a Student’s t-test, p<0.001,t=4.60,df=13, n=6-9/group, Fig. 5*B*; Mean pT205 immunopositive area fractions in the CA1: 0.01 ± 0.002 for SC560 treated mice vs. 0.14 ± 0.04 for vehicle treated mice; Data analyzed by a Student’s t-test, p<0.01,t=3.65,df=13, n=6-9/group, Fig. 5*C*). In comparison pT205 immunopositive area fractions between SC560 and vehicle treated Tau mice were not significantly different from each other (data not shown). To validate the phospho-tau data, we conducted western blot analysis of hippocampal lysates from vehicle or SC560 treated mice. Vehicle treated Tau/IL-1 mice showed a prominent upregulation of phosphorylation at the pT205 and PHF1 epitopes. SC560 treatment led to significantly reduced phosphorylation at both these epitopes in the Tau/IL-1 mice. SC560 and vehicle treated Tau mice demonstrated similar levels of phosphorylation at pT205 and PHF1 epitopes (Fig. 5*D*). Densitometric quantification of band intensities revealed a 2-3 fold induction of phosphorylation at the pT205 and PHF1 epitopes following IL-1β overexpression, when normalized to total-tau levels (Mean densitometric values for pT205/total-tau in arbitrary units: 0.2674 ± 0.05 for vehicle treated Tau mice vs. 0.822 ± 0.08 pg/mg protein for vehicle treated Tau/IL-1 mice; Mean densitometric values for PHF1/total-tau in arbitrary units: 0.382 ± 0.02 for vehicle treated Tau mice vs. 0.883 ± 0.02 for vehicle treated Tau/IL-1 mice). SC560 treatment reduced tau phosphorylation at both these epitopes (Mean densitometric values for pT205/total-tau in arbitrary units: 0.554 ± 0.06 for SC560 treated Tau/IL-1 mice vs. 0.822 ± 0.08 for vehicle treated Tau/IL-1 mice; Mean densitometric values for PHF1/total-tau in arbitrary units: 0.702 ± 0.02 for SC560 treated Tau/IL-1 mice vs. 0.883 ± 0.08 for vehicle treated Tau/IL-1 mice; n=4-6/group; Data analyzed by a one way ANOVA followed by a Tukey’s post-hoc test for individual comparisons; p<0.001; Fig 5*E,F*).

**Fig. 5.**
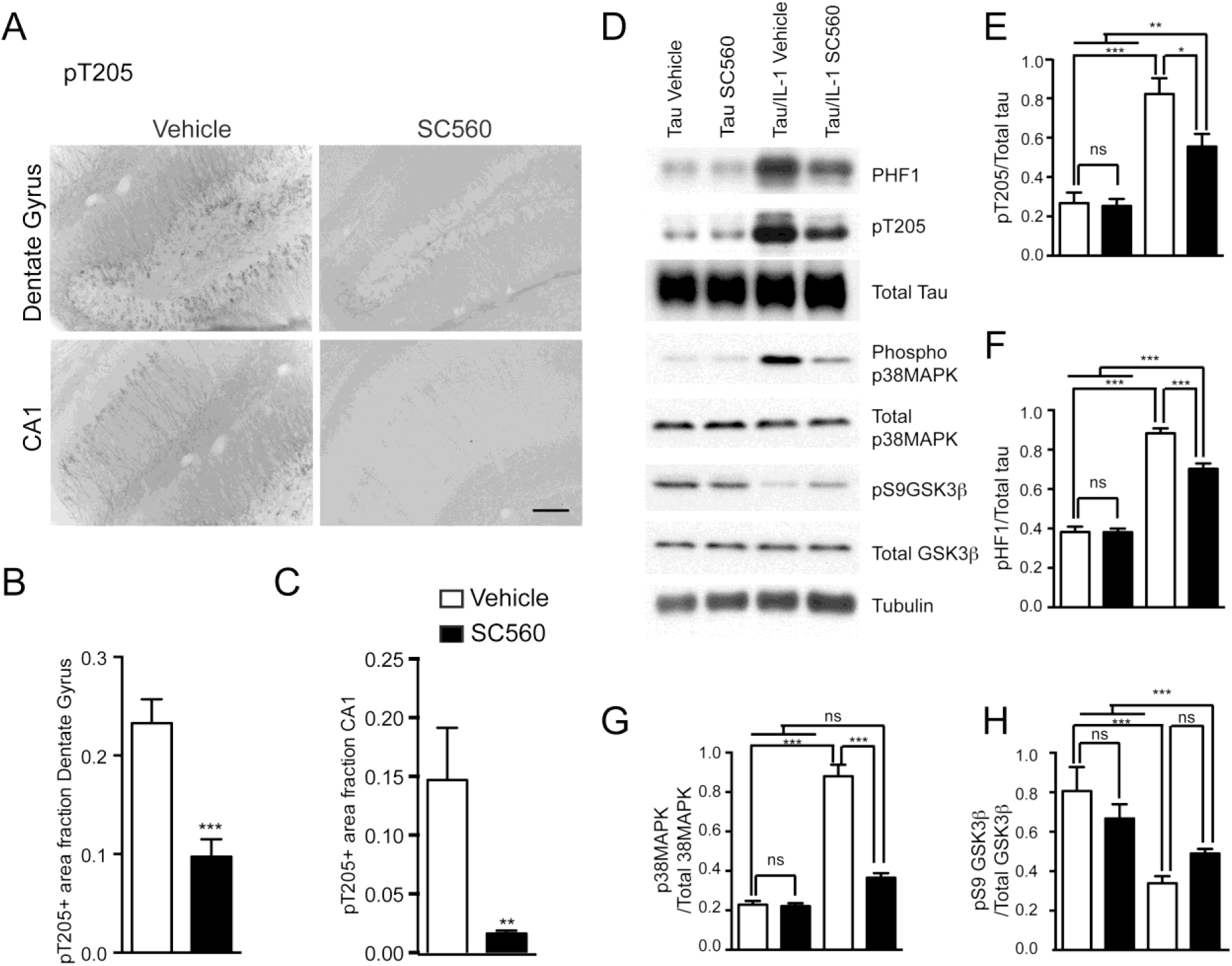
SC560 treatment reduces tau pathology and active tau kinases in 9-month-old Tau/IL-1 mice. Representative micrographs of sections from the dentate gyrus and CA1 of 9-month-old Tau/IL-1 mice immunostained for phospho-tau (pT205) are shown. One week of treatment with SC560 reduced pT205 immunostaining both in the dentate and CA1 of Tau/IL-1 mice despite overexpression of the IL-1β transgene for 4 weeks ***(A)***; Scale bars 100 μm. Quantification of pT205 immunopositive area fraction in the dentate gyrus ***(B)*** and CA1 ***(C)*** of the hippocampus of Tau/IL-1 mice showed significant reduction in SC560 treated groups (black bars) compared to vehicle treated groups (open bars). All data are represented as mean pT205 immunopositive area fraction ± SEM per group. Data analyzed by a Student’s t-test. n=6-9/group. *** p<0.001; ** p<0.01. ***(D)*** Representative western blots probed with various antibodies against phosphorylated and total tau and tau kinases are shown. Overexpression of the IL-1β transgene for 1 month led to significant upregulation of phosphorylation at pT205 and PHF1 epitopes, which was mitigated by SC560 treatment for 1 week ***(D, E, F)***. IL-1β overexpression also increased the phosphorylation of p38MAPK, which was again reversed by SC560 treatment ***(D, G)***. Further, IL-1β overexpression reduced phosphorylation at Ser9 of GSK3β, which showed a trend of increase following SC560 treatment ***(D, H)***. The steady-state levels of Tau, p38MAPK and GSK3β were not altered between genotypes across treatments. For detailed description of quantitative outcomes please refer to the text. All data are expressed as mean densitometric counts in arbitrary units of phospho/total proteins ± SEM per group. Data analyzed by a one-way ANOVA with Tukey’s multiple comparison post-test for individual comparisons. n= 4-8/group. *** p<0.001; * p<0.05; ns: not significant

### 3.5. Pharmacological inhibition of COX-1 by SC560 reduces active forms of tau kinases p38MAPK and GSK3β which are induced in the Tau/IL-1 mice following IL-1β overexpression

We investigated if SC560 treatment could modulate changes in potential tau kinases. Previously, we reported upregulation in active p38MAPK and disinhibition of GSK3β in triple transgenic Alzheimer’s mice following IL-1β overexpression (Ghosh et al. 2013). In agreement with that data, we observed significant upregulation in phospho-p38MAPK, suggesting increased activity of p38MAPK in the Tau/IL-1 mice that received vehicle, compared to vehicle treated Tau mice (Fig. 5*D,G*). Additionally, Tau/IL-1 mice demonstrated significantly less phosphorylation at the Ser9 epitope (pS9) of GSK3β, which is an inhibitory phosphorylation for GSK3β. A decrease in this epitope reflects a withdrawal of inhibition, and therefore increased GSK3β activity in the Tau/IL-1 mice (Fig. 5*D,H*). SC560 treated Tau/IL-1 mice demonstrate a significant reduction in phospho-p38MAPK, and a trend towards increased pS9GSK3β None of the kinases demonstrated any change in the total levels (Fig. 5*D,G-H*; Mean densitometric values for phospho-p38MAPK/total p38MAPK in arbitrary units: 0.228 ± 0.01 for vehicle treated Tau mice vs. 0.880 ± 0.05 for vehicle treated Tau/IL-1 mice; Mean densitometric values for pS9GSK3β/total GSK3β in arbitrary units: 0.806 ± 0.16 for vehicle treated Tau mice vs. 0.339 ± 0.03 for vehicle treated Tau/IL-1 mice; Mean densitometric values for phospho-p38MAPK/total p38MAPK and pS9GSK3β/total GSK3β in arbitrary units for SC560 treated Tau/IL-1 mice: 0.365 ± 0.02 and 0.489 ± 0.02 respectively; n=4-6/group; Data analyzed by a one-way ANOVA with Tukey’s post-hoc test for individual comparisons; p<0.001).

## 4. Discussion

Tauopathies consist of a spectrum of disorders related to mutations in the tau gene (Ballatore et al. 2007). Frontotemporal dementia (FTD), the second most common form of tauopathies following Alzheimer’s disease, accounts for 5-10% of diagnosed dementia cases overall, and is the third most common form of dementia across all age groups (Seltman and Matthews 2012). FTDs typically strike earlier than AD, with 60% of cases being detected between 45-65 years of age (Knopman and Roberts 2011). The time from onset of symptoms to death is estimated to be between 6.5-11 years (Seltman and Matthews 2012), during which patients steadily deteriorate in cognitive, executive and motor performance, thus becoming increasingly dependent on caregivers (Boxer et al. 2013a). Currently, there is no FDA approved disease modifying agent to treat FTD, although SSRIs, cholinesterase inhibitors, and NMDA channel blockers are used for symptomatic treatment (Boxer et al. 2013b). Therefore, it is crucial to identify and pursue therapeutic targets. Along with tau inclusions, neuroinflammation figures prominently in FTD pathogenesis. The FTD brain shows increased gliosis and inflammatory cytokines and mediators (Boxer et al. 2013a). In a transgenic mouse model of FTD, microglial activation preceded neurofibrillary pathology and degeneration (Yoshiyama et al. 2007). Along those lines prominent microglial activation is present in the FTD brain as detected by PET (Cagnin et al. 2004), and parallels disease pathogenesis in PSP and CBD brains (Ishizawa and Dickson 2001). In P301S transgenic mice demonstrating abundant filamentous tau pathology (Allen et al. 2002), IL-1β and COX-2 immunoreactivity is markedly elevated in brainstem and spinal cord neurons, and microglia are prominently immunoreactive for IL-1β (Bellucci et al. 2004). Preliminary studies also showed robust microglial CD68 immunoreactivity in close proximity of phospho tau positive inclusions in an FTD patient, suggesting close association of phagocytic microglia with pathological tau inclusions. In the same brain, IL-1β and COX-2 colocalized with phospho-tau markers such as AT8 and AT100 (Bellucci et al. 2011). We sought to characterize the relationship of tau pathology and IL-1β in further detail and used the well-characterized IL-1β^XAT^ mouse developed in our laboratory (Shaftel et al. 2007b).

In this study we present direct evidence that sustained upregulation of IL-1β exacerbates tau phosphorylation in a tau transgenic model. Our data is supported by recent observations in several models suggesting immune activation *in vivo* can exacerbate tau pathology. For example, central LPS injections worsen tau pathology in the rTg4510 model of forebrain specific transgenic tau overexpression (Lee et al. 2010). Moreover, in a model overexpressing wild-type human tau, ablating the microglial Fractalkine receptor (CX3CR1) resulted in more aggressive glial activation and exacerbated phospho-tau pathology. This effect was demonstrated to be dependent on IL-1 and TLR4 signaling (Bhaskar et al. 2010). Further, indirectly blocking IL-1 signaling by peripheral administration of an antibody against IL-1R1, the principal IL-1 receptor attenuates tau phosphorylation in 3xTgAD mice (Kitazawa et al. 2011). These data are consistent with our current results, and our previous observation of an IL-1β-mediated exacerbation of tau pathology in 3xTgAD mice (Ghosh et al. 2013). We observed markedly elevated microglial activation in Tau/IL-1 mice at both time points assessed in our study, suggesting a possible role of microglia in the observed elevation of phospho-tau. We also obtained evidence suggesting increased activity of p38MAPK and possibly GSK3β in Tau/IL-1 mice. p38MAPK colocalizes with AT8 positive and NFT bearing neurons in human AD brain tissue (Zhu et al. 2000;Ferrer et al. 2001;Sheng et al. 2001), which underscores the importance of this kinase in tau phosphorylation. IL-1β treatment exacerbates tau phosphorylation at the AT8 epitope, which is mediated by p38MAPK in primary rat cortical neurons (Li et al. 2003a). Rat brains impregnated with slow releasing pellets of IL-1 show upregulation in p38MAPK mRNA (Sheng et al. 2001). In the current study with JNPL3 mice, we observed increased phospho-p38MAPK in hippocampal lysates with IL-1β overexpression, which provides a direct link between IL-1β and tau phosphorylation. Additionally, we demonstrated a disinhibition of GSK3β with IL-1β in Tau/IL-1 mice, which has been extensively characterized as a relevant tau kinase in AD (Jope and Johnson 2004;Kremer et al. 2011). Whether upregulation of these kinases are a direct effect of IL-1β overexpression cannot be ascertained without further mechanistic experiments. However, our work suggests that both these kinases are important regulators of tau pathology in the context of inflammation.

Of note, our model elicits a focal inflammatory response that is largely restricted within the hippocampus in experimental animals, and therefore, this experiment provides proof of concept type data following IL-1β overexpression in a location specific manner, in an animal that expresses transgenic human tau. The slow global neuroinflammatory response in human tauopathies are not truly mimicked by any existing mouse model, however, we feel that within the normal lifespan of a tau transgenic mouse (Often around thirteen to fourteen months), a one to three month period of overexpression is reasonably long term and can be comfortably utilized for meaningful interpretation of data.

The second major observation of this study is that pharmacological inhibition of COX-1 by the selective inhibitor SC560 reduced the neuroinflammatory mediators mIL-1β and PGE_2_, decreased microglial activation and attenuated phospho-tau pathology in Tau/IL-1 mice compared to vehicle treated mice. Additionally, we demonstrated that increased active p38MAPK in vehicle treated Tau/IL-1 mice was reduced back to control levels in SC560 treated Tau/IL-1 animals. Active GSK3β also tended to decrease with SC560 treatment, although the effect did not quite reach significance. PGE_2_ can directly compromise hippocampal learning in rodents (Hein and O’Banion 2009); therefore, its reduction in the Tau/IL-1 mice may have important behavioral benefits that might be addressed in a future experiment. Notably, we did not observe any change in phospho-tau pathology or microglial activation in mice lacking the IL-1β transgene with SC560 treatment. This contrasts with recent data from an experiment with 3xTgAD mice in which SC560 treatment led to attenuation of both amyloid and tau pathology with corresponding functional improvement (Choi et al. 2013). However, this study used 20-month-old 3xTgAD mice, which demonstrate advanced amyloid pathology, and a proportionally strong neuroinflammatory response, including prominent microglial activation. Amyloid is a well-known activating stimulus for microglia, which the Tau mice lack. In contrast to the 3xTgAD mice, which show extensive microglial activation (Kitazawa et al. 2004), the 8 month old Tau mice used for this study show little or no phenotypic activation of microglia (data not shown). It might be possible that the COX-1 pathway is not sufficiently active in the Tau mice for SC560 to make a big difference in pathological outcomes at the time points chosen. However, once the IL-1β transgene is activated in our model and escalates microglial activation and the overall neuroinflammatory response, COX-1 appears to become an important player in regulating tau pathology. Further experiments will be needed to elucidate underlying mechanisms of these effects. For example, inhibiting SC560 in a more aggressive model of tauopathy like the P301S transgenic mouse, which has a strong inflammatory component (Allen et al. 2002;Yoshiyama et al. 2007), might provide rescue from pathology and functional deficits.

Considerable literature implicates the involvement of COX pathways in chronic neurodegenerative disorders like AD (Heneka and O’Banion 2007;Heneka et al. 2010). Several lines of evidence suggest that COX-1 inhibition might be a better therapeutic strategy to attenuate neuroinflammation in chronic neurodegeneration than COX-2 inhibition. Unlike COX-2, which is predominantly expressed in neurons in the CNS (O’Banion 1999), COX-1 is expressed in microglia in rodent and human brains and is slightly elevated in AD (Yermakova et al. 1999). Therefore, the localization of COX-1 is consistent with it responding to acute or chronic inflammatory stimuli (Aid and Bosetti 2011). COX-1 has also been implicated in PGE_2_ synthesis in a number of CNS injury models (Heneka et al. 2010), while COX2 has been implicated in a range of anti-inflammatory functions (Aid and Bosetti 2011). Interestingly, COX-2 inhibition was associated with exacerbated amyloid pathology in clinical trials, while COX-1 inhibition was associated with slightly better outcomes (McGeer and McGeer 2013). COX-1 inhibition has been used as a therapeutic strategy before to attenuate neuroinflammatory responses in stroke models (Candelario-Jalil et al. 2003) and COX-1 knockout mice have decreased inflammatory responses, oxidative stress and neuronal damage following central LPS or Aβ42 challenge (Choi et al. 2008;Choi and Bosetti 2009). Finally, as already mentioned, SC560 improved pathological outcomes in 3xTgAD mice (Choi et al. 2013). Therefore, our observations in the Tau mice add to a growing body of evidence that COX-1 is likely to be an important driver of neuroinflammatory responses in chronic neurodegenerative disorders. Following the apparent failure of NSAIDs in clinical trials of AD patients, interest in these drugs as therapeutic agents has declined, however, when taken together, the preclinical data in this study and that of others suggests that a careful reevaluation is essential, especially with regard to COX-1 specific inhibitors.

In summary, we have established that IL-β upregulation exacerbates tau pathology in Tau transgenic mice in a COX-1 dependent manner. This suggests that inhibition of COX-1 might be a valuable therapeutic strategy in human tauopathies with an advanced inflammatory component.

## Disclosure statement

The authors disclose no conflicts of interest.

## Acknowledgements

This work was supported by a grant from the National Institute of Aging (RO1 AG30149). The authors thank Jen-nie Miller for preparation of the FIV-Cre virus stock, and Jack R. Walter, Lee A.Trojanczyk and Mallory E. Seaman for assistance with genotyping, histology and animal colony maintenance. The PHF1 antibody used in this study was a generous gift of Dr. Peter Davies, Albert Einstein College of Medicine, New York, NY.

---

**Fig. S1.**
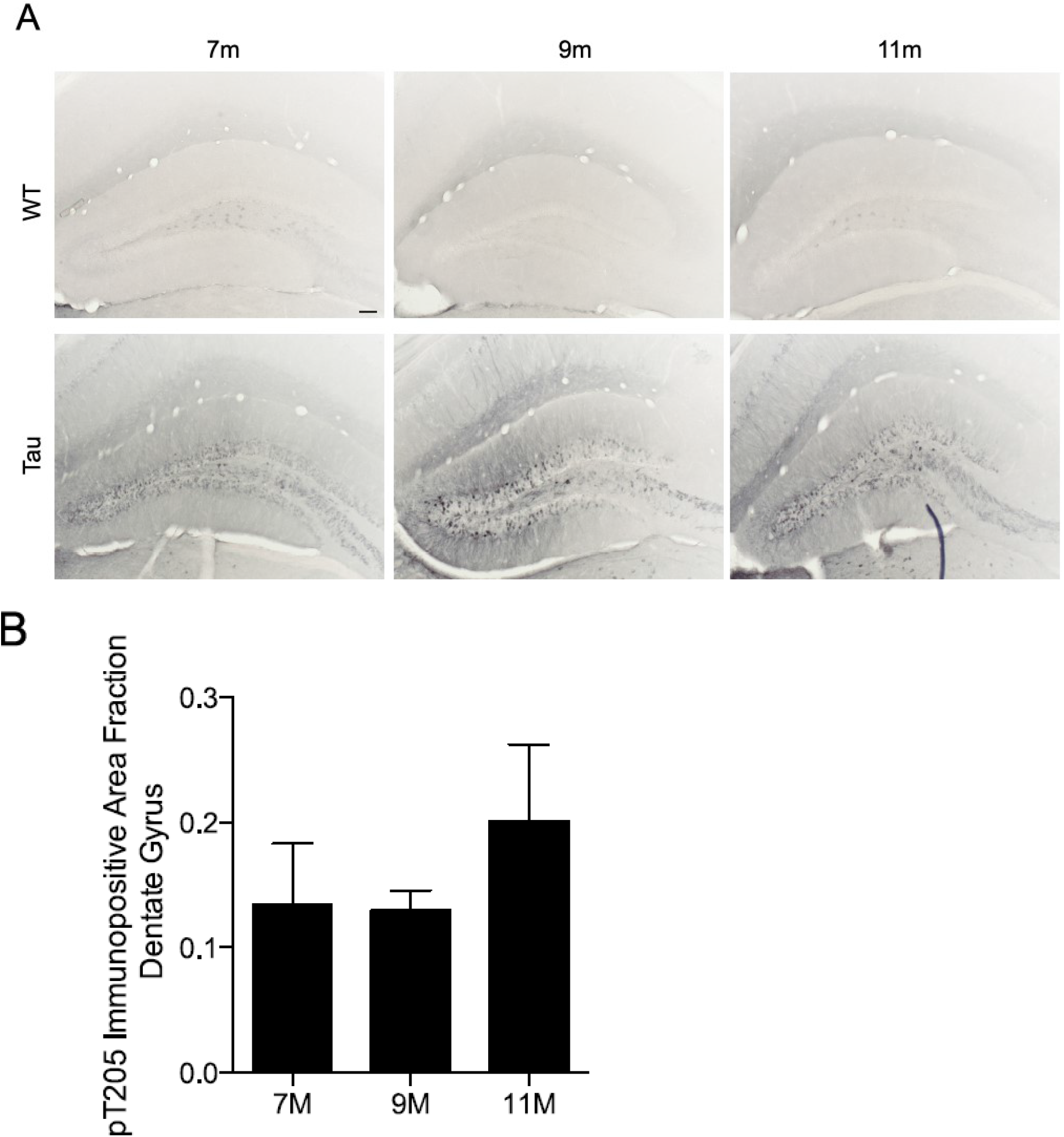
pT205 immunostaining is detectable in Tau mice at 7, 9 and 11months of age but not in littermate WT controls. Representative photomicrographs from dentate gyrus of Tau and WT mice immunostained for pT205, a pathological phospho-Tau epitope are shown (***A***) Scale bars 100 μm. ***(B)*** Quantification of pT205 immunopositive area fractions are shown (n=3-5 animals/group)

**Fig. S2.**
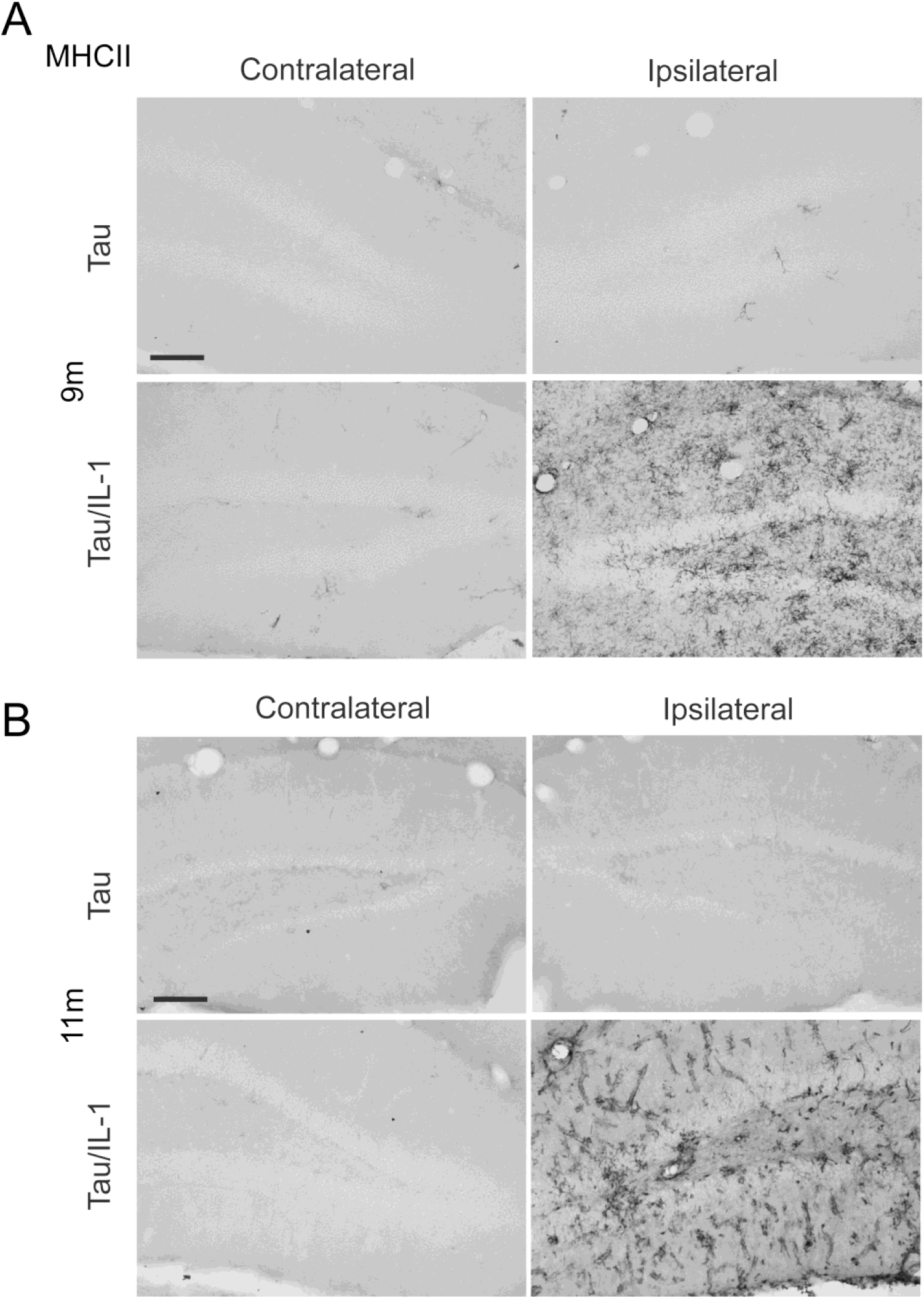
MHCII is upregulated in Tau/IL-1 mice following one and three months of IL-1β overexpression. Representative photomicrographs from dentate gyrus of Tau and Tau/IL-1 mice immunostained for the inflammatory marker MHCII are shown. MHCII was upregulated in the ipsilateral dentate gyrus following overexpression of the IL-1β transgene for one (***A***) and three (***B***) month in 9 month and 11 month old Tau/IL-1 mice respectively. Control Tau mice lacking the IL-1β transgene at these same ages did not show any significant MHCII staining in either of the hemispheres. Scale bars 100 μm.

**Fig. S3.**
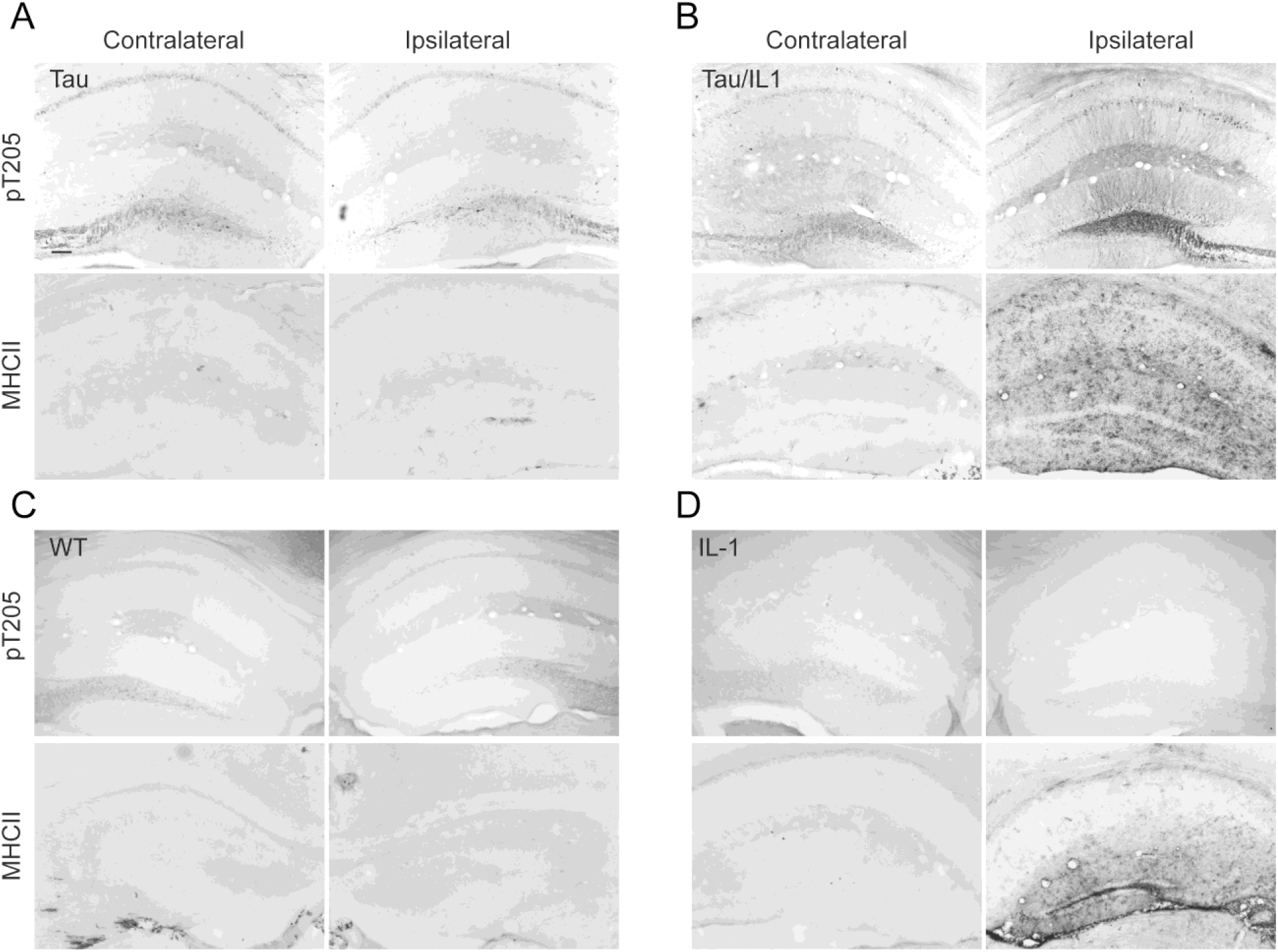
MHCII and pT205 immunostaining at 9 months. All four genotypes i.e. WT, IL-1, Tau and Tau-IL-1 were immunostained for an inflammatory marker (MHCII) and phospho-tau (pT205). MHCII was upregulated in the ipsilateral hippocampus of 9 month old Tau/IL-1 and IL-1 mice ***(B, D)*** but not in Tau or WT mice ***(A, C)*** which do not have the IL-1β transgene. pT205 immunostaining was detected in Tau and Tau/IL-1 mice ***(A, B)***, but not in WT or IL-1 mice ***(C, D)***, which do not express transgenic tau. pT205 immunostaining was increased in the ipsilateral hippocampus of Tau/IL-1 mice following one month of IL-1β overexpression. Scale bar 100 μm.

**Fig. S4.**
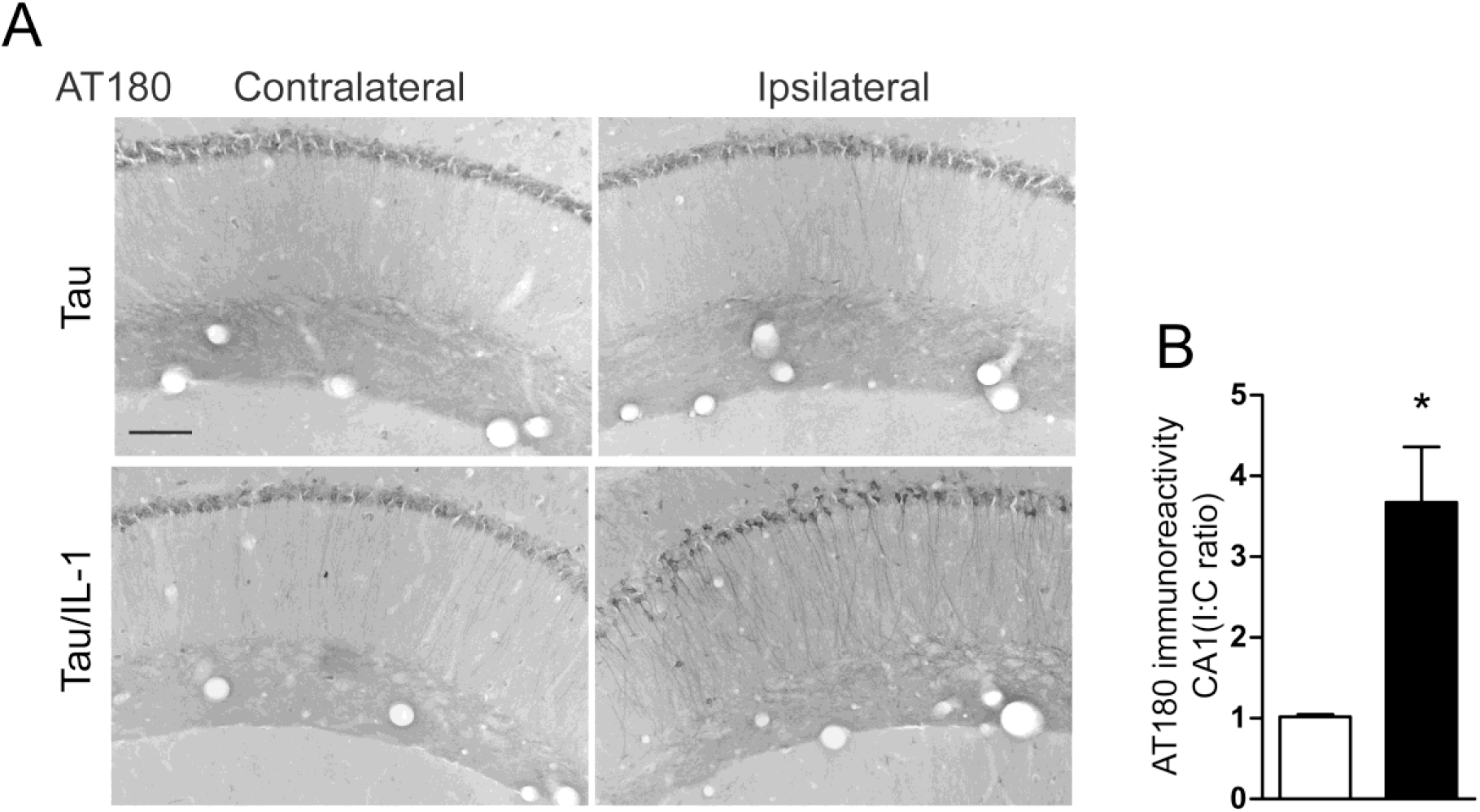
9 month old Tau/IL-1 mice show increased AT180 immunostaining in the ipsilateral CA1 area. Sections of the hippocampus of were probed with antibodies against phospho-tau (pT205 and AT180). Representative images of hippocampal sections from 9 month old Tau and Tau/IL-1 mice stained with AT180 (***A***) show increased immunostaining in the ipsilateral CA1 regions of Tau/IL-1 mice at 9 months of age (after one month of IL-1β overexpression). Scale bar 100 μm. ***(B)*** One month of IL-1β overexpression significantly upregulates phospho-tau at the AT180 epitope in the ipsilateral CA1 of Tau/IL-1 mice. n= 5-6/group, Student’s t-test, *p<0.05.

## Notes

### Competing Interest Statement

The authors have declared no competing interest.

